# Disturbance provides limited respite for native fish against invaders

**DOI:** 10.1101/2025.08.22.671881

**Authors:** Julian Merder, Angus R. McIntosh, Jan A. Freund, Rory S. Lennox, Naomi Heller, Jonathan D. Tonkin

## Abstract

Understanding how flow-related disturbance regimes influence species interactions is critical for conserving threatened species in freshwater ecosystems, where both the alteration of these regimes and the invasion of non-native species pose major threats. Using 30 years of sampling data from a well-studied river system in Aotearoa New Zealand, we applied empirically– parameterized Lotka–Volterra competition models across a gradient of flood disturbance regimes to assess the potential for coexistence between threatened native galaxiids and invasive trout. Our models suggest trout dominate under low disturbance regimes, leading to galaxiid extinction. Coexistence peaks at intermediate disturbance regimes, though native fish densities are highly suppressed. Only at high disturbance regimes do native fish prevail, but priority effects are likely. Thus, restoration and conservation efforts focused solely on high disturbance habitats without accompanying biological interventions may not ensure native fish persistence. Further, prioritizing such habitats at the expense of others risks misdirecting resources towards sub-optimal conditions, particularly under climate change.

## Introduction

Freshwater ecosystems face increasing threats from invasive species, reflecting their role as a central driver of global biodiversity loss (IPBES 2023). Invasive freshwater fish, in particular, have caused widespread ecological and economic impacts, with global costs exceeding $37 billion since 1960 (Haubrock *et al*. 2022). More than 550 alien freshwater fish species have already established populations outside their native basins (Bernery *et al*. 2022), with introductions continuing at an accelerating pace. Invasions by larger-bodied fish, including carp and trout, now affect about 59% of the world’s primary river basins (Chen *et al*. 2024).

Critical to the spread of invasive freshwater species in much of the world has been the modification of river flow regimes (Bunn and Arthington 2002). These regimes, including floods and droughts, are fundamental disturbance forces driving the evolution of life histories, population dynamics, and community assembly (Bunn and Arthington 2002; Lytle and Poff 2004). Thus, a key tool in managing the impacts of invasive species in rivers has been targeting components of the natural disturbance regime in flow reoperation (Chen and Olden 2017; Tonkin *et al*. 2021). However, this approach rests upon the assumption that native species, adapted to the natural flow regime (Lytle and Poff 2004), gain a competitive advantage from natural disturbances, providing a buffer against invaders. Realistically, however, the potential for natural disturbance regimes to protect native fishes from invasion is likely highly context dependent, given that the natural regime itself is dynamic, ranging from minimally disturbed to uninhabitable at the landscape scale (McIntosh and Barrett 2022). Habitats at the upper end of this gradient can become too harsh even for native species, decreasing population growth and reproduction, negative effects that need to be outweighed by potentially lowered competition (Chesson and Huntly 1997; McIntosh and Barrett 2022).

Understanding how this trade-off between invasibility and disturbance tolerance unfolds across disturbance gradients is essential for effective conservation efforts. This issue is particularly pertinent in the context of climate change, which is anticipated to increase the frequency and magnitude of floods and droughts (Tonkin *et al*. 2019), risking native species falling into disturbance-driven ecological traps (Battin 2004). Given the highly contested arena within which water management decisions operate, including trade-offs among competing human and ecosystem demands (Chen and Olden 2017) and among sectors of the ecosystem (Tonkin *et al*. 2021), it is essential to base conservation decisions on sound mechanisms.

In Aotearoa New Zealand, more than 70% of galaxiid fish species are threatened with extinction, and invasive brown (*Salmo trutta*) and rainbow trout (*Oncorhynchus mykiss*) are widely regarded as a major cause of their decline (McDowall 2006). River resident galaxiids—a major component of the endemic freshwater fish fauna that are ecologically and culturally integral to Aotearoa—are particularly threatened. While previous research suggests that these galaxiids fare better in more disturbed streams (Leprieur *et al*. 2006), such conditions are likely suboptimal, serving as present-day refuges from trout competition (Hore *et al*. 2025). Conservation strategies could mistakenly rely on promoting these high disturbance regimes, not recognizing the risks they pose, especially as conditions become harsher. To test whether and how disturbance regimes mediate the coexistence or dominance of native and non-native fish, we applied modern coexistence theory to a 30-year fish community dataset to quantify the competitive outcomes between native and invasive fishes across a disturbance gradient from stable to highly disturbed (Figure 1). Our approach provides critical insights into when and how natural disturbance regimes can promote the persistence of native species experiencing invasion, and the potential for perverse outcomes from management focused solely on disturbance regimes, ultimately informing more effective conservation strategies both locally and globally.

**Figure 1:**
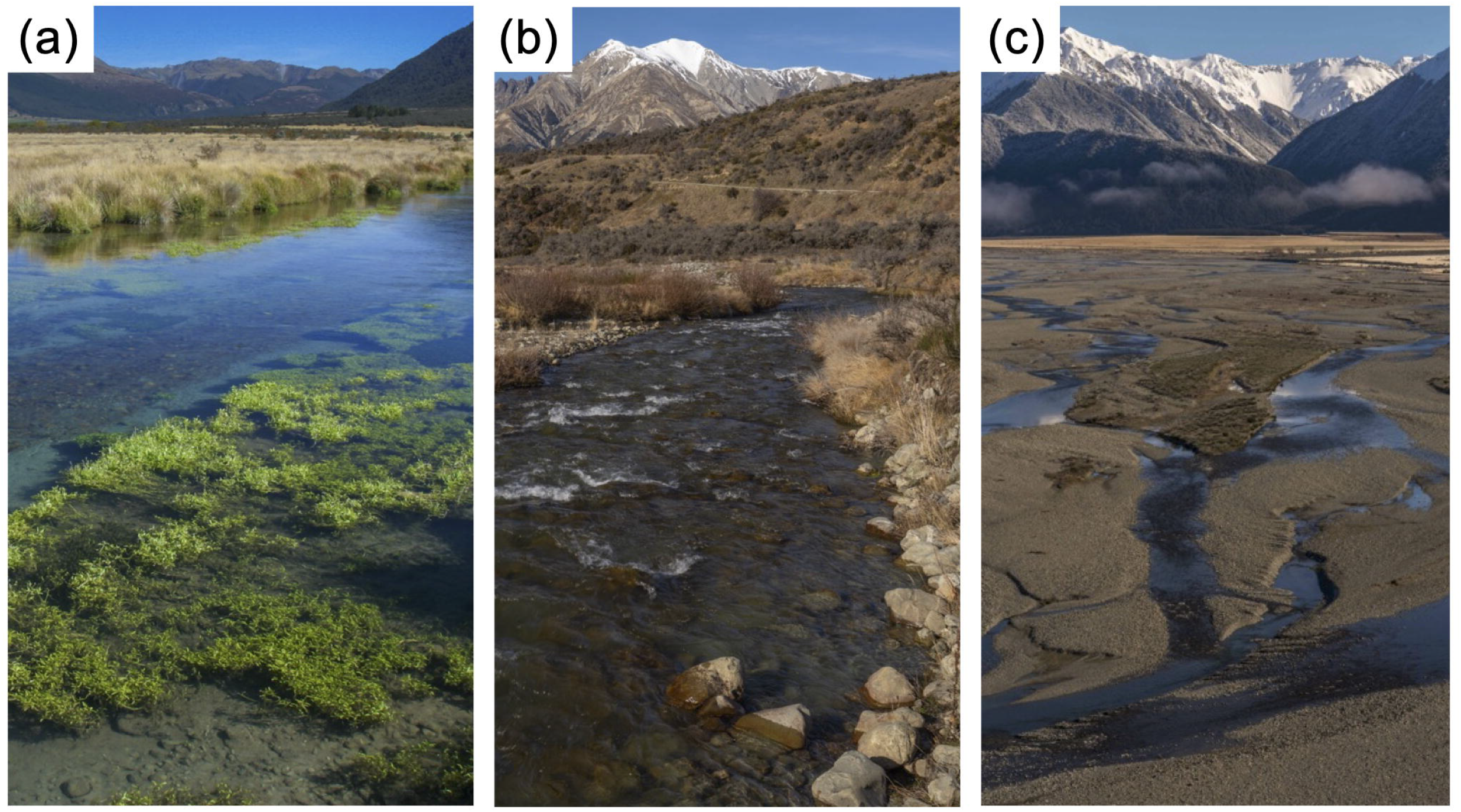
Examples of sites within the study catchment (Waimakariri River, New Zealand) representing the three disturbance regimes: (a) low, (b) intermediate, and (c) high disturbance. Image credits: *Angus R. McIntosh*

## Methods

We used a dataset of 10,243 fish catch records collected over 30 years across 424 fishing occasions in the upper Waimakariri River catchment. Each catch, classified as either trout (*Salmo trutta, Oncorhynchus mykiss*) or galaxiid (*Galaxias vulgaris, G. paucispondylus*), is paired with a Pfankuch Index value—a standardized measure of streambed disturbance, found to better reflect responses in fish biomass and assemblage structure than purely hydrological measures (Jellyman *et al*. 2013). We categorized disturbance regimes into three levels (Figure 1): low (≤70), medium (71–100), and high (>100). For the analysis, we treated the two galaxiid species as a single river resident galaxiid guild (G) despite differences in early life history (Jones and Closs 2016), due to the general consistency of responses of adults to disturbance and trout (Hore *et al*. 2025). Similarly, we treated the two trout species as a single guild (S) because of their consistent life histories and morphology in these small high-elevation streams (typical maximum size ∽250 mm).

To mechanistically understand how the trade-off between invasibility and disturbance tolerance shapes competitive outcomes across natural disturbance regimes, we modeled population dynamics using Lotka-Volterra type competition equations (Chesson and Huntly 1997; Chesson and Kuang 2008), incorporating maximal *per-capita* growth rates (*r*_*G*_,*r*_*S*_), interspecific competition coefficients (*a*_*GS*_, *a*_*SG*_) and intraspecific competition coefficients (*a*_*GG*_, *a*_*SS*_) (Appendix S1: Panel S1). While the data are extensive, times between sampling events are not equidistant and vary from years to decades. As such, we used space-for-time substitution by looking at the relationship between total and adult fish densities (*G*_*A*_,*S*_*A*_) at a given location as an approximation for the change between densities over time (Appendix S1: Panel S1). Since both trout and galaxiids have relatively short lifespans—whose juvenile stage typically lasts only one year—this approach captures a transition roughly equivalent to the shift from one cohort to the subsequent adult stage.

We then estimated parameters using Bayesian regression, excluding samples where and densities were zero to avoid natural logarithms used in Lotka-Volterra equations becoming undefined. We fitted parameters simultaneously in a combined brms model (Appendix S1: Panel S2) where coefficients were estimated for each level of disturbance (low, medium, high) in the brms (v. 2.22.0) package in R (Bürkner 2017). Differences in densities due to season were included as cyclic smooth functions of month for the growth rate parameters to account for seasonal heterogeneity, which was treated as a random effect within brms (Bürkner 2017). We used a lognormal distribution as a conditional distribution for fish densities.

Based on the fitted Lotka-Volterra model parameters, we used methods from modern coexistence theory (Chesson and Kuang 2008) to estimate whether trout could invade and whether coexistence occurred, or if trout drove galaxiids to extinction, by calculating niche differences and fitness ratios for each disturbance regime. Niche difference (ND) measures the extent to which within-species competition is stronger than between-species competition, indicating whether species are more limited by themselves or by other species. The fitness ratio (FR) reflects their relative competitive abilities, meaning that when FR <1, galaxiids have a fitness advantage over trout. Coexistence occurs when niche differences are significant enough to offset fitness differences (Appendix S1: Panel S3). If niche difference is negative, indicating that species compete more strongly with each other than with themselves, interactions become destabilizing and can lead to priority effects, where the outcome depends on initial abundances. This implies that if enough galaxiids are present, trout cannot invade. However, if galaxiid populations are very low and many trout arrive, dynamics may tip toward a trout-only system.

We visualized the results of estimated model parameters using phase plots with equilibrium outcomes highlighted. Our Bayesian model fits also allowed further incorporation of parameter uncertainties in coexistence predictions. To achieve this, we drew 10,000 samples from the posterior distribution of coefficients, recalculating the fitness ratio and niche difference, and determining whether coexistence was expected for each disturbance regime. Furthermore, we calculated the proportion of draws that yielded either coexistence, priority effects, trout dominance, or galaxiid dominance, thus reflecting the sensitivity of the overall pattern and its robustness (Terry and Armitage 2024).

## Results and discussion

We found a clear ordering of trout and galaxiid competitive hierarchies along the disturbance regime gradient (Figure 2a). At low disturbance, trout overwhelmingly dominated the community, with an 82% probability of driving galaxiids extinct (Figure 2b). Fitness ratios were clearly in favor of trout (Figure 2b), with the chance of galaxiid dominance approaching zero (including the lowest associated growth rates across regimes), and a low probability of coexistence (11%). The low growth rates of galaxiid populations, alongside high galaxiid–trout niche differences at low disturbance, may reflect the greater abundance of protected consumers (e.g., cased or shelled invertebrates) found in spring-fed streams in this region, which trout exploit more effectively than galaxiids (Jellyman and McIntosh 2020). This pattern is consistent with the high trout growth rates and highest carrying capacities (1/*a*_*ss*_) observed at low disturbance rates (Appendix S1: Figure S1a). These results align with a recent meta-analysis that shows trout prefer habitats with stable, mid-velocity conditions suitable for drift feeding (Rosenfeld and Enright 2025).

**Figure 2:**
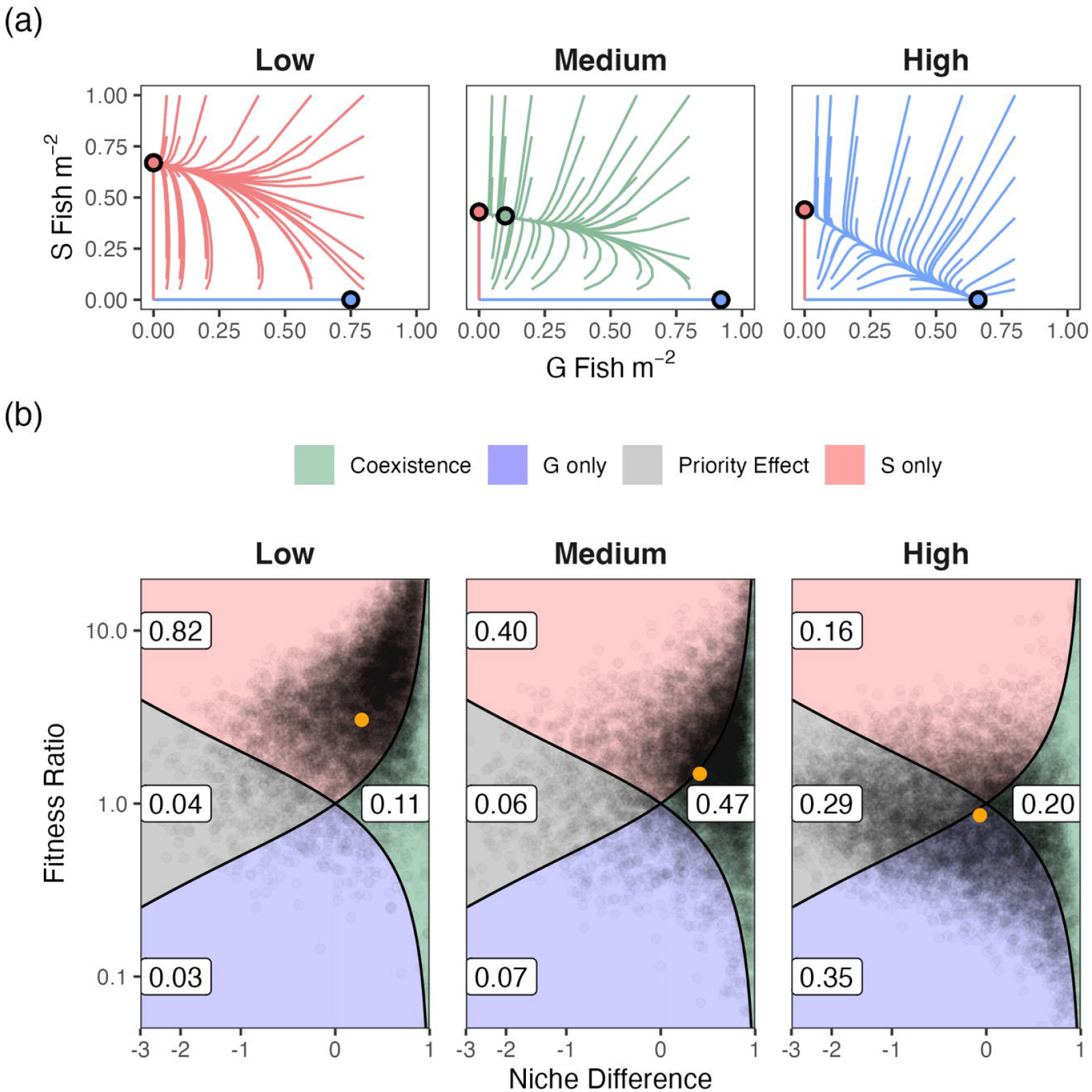
Predictive competitive outcomes between galaxiids (G) and trout (S) are shown as (a) phase plots illustrating dynamic trajectories under varying disturbance levels, based on mean parameter estimates, and (b) parameter space of niche differences and fitness ratios capturing outcome uncertainty from 10,000 posterior draws of our Bayesian regression model. In (a), each panel shows species density trajectories from varied initial conditions, with colored lines indicating direction and circles marking equilibrium points (stable or unstable depending on flow field color). In (b), shaded regions represent competitive outcomes, black circles posterior draws (with percentages), and orange circles indicate mean estimates shown in (a).

By contrast, at high disturbance levels, the risk of disruptive trout invasion was minimal (16%) and the probability of coexistence was higher (20%) compared to the low disturbance regime. Under these conditions, galaxiids gained a slight competitive advantage, with their probability of outcompeting trout maximized (35%)—still modest and highly disproportionate compared to the dominance of trout at low disturbance (85%). This reduced invasion risk suggests that trout are less well adapted to such harsh conditions, supporting findings by Fausch *et al*. (2001), who showed that flood disturbance limits the invasion success of rainbow trout across five Holarctic regions—particularly in systems with frequent, unpredictable flooding. Under such high disturbance conditions, the risk of salmonid eggs and fry being destroyed or displaced is high. Consistent with these challenging conditions, estimated carrying capacities under high disturbance regimes were lower for both trout and galaxiids (1/*a*_*ss*_, 1/*a*_*GG*_) compared to the low disturbance regime, emphasizing that harsh environments tend to generally support lower species densities (Appendix S1: Figure S1a). However, while trout exhibited their lowest growth rates here, growth rates remained higher than those of galaxiids (Appendix S1: Figure S1a).

Importantly, the probability of priority effects increased along the disturbance regime gradient, becoming very likely at high disturbance levels (29%). Although this result contrasts with experiments suggesting priority effects are less likely under harsh conditions (droughts) in aquatic communities (Chase 2007), it highlights that harshness is not necessarily a relief from competition (Chesson and Huntly 1997). Fitness ratios and niche differences converging to parity, leading to such priority effects, indicate that stochastic processes play an increasingly central role in determining the persistence of native species in harsh conditions (Hubbell 2006), which can modulate the success of conservation measures. Since native species are unable to sufficiently outcompete trout at either end of the disturbance spectrum, their persistence becomes even less likely under conditions that exceed the boundaries of the natural regime. Previous empirical work shows that anthropogenically altered flow regimes and climate-induced reduced streamflow favor invaders (Ruhí *et al*. 2016; Rogosch *et al*. 2019). Our results further support the concern that conservation strategies promoting regimes near the upper end of natural disturbance may act as ecological traps (Battin 2004)—particularly as climate change intensifies environmental harshness, placing additional pressure on already struggling native populations.

The highest probability of coexistence (47%) occurred at intermediate disturbance regimes. Intermediate disturbance rates do not, however, guarantee native fish persistence, because our model also predicted that trout winning remained more likely than galaxiids outcompeting trout. Moreover, the mean model parameters indicated very low estimated galaxiid densities at coexistence equilibrium (Figure 2a), further emphasizing the imbalanced outcomes in favor of trout, which show equilibrium densities close to their carrying capacities (Figure 2a). The suppression of galaxiid densities is particularly severe given that, at intermediate disturbance regimes, galaxiids exhibit their peak growth rates and carrying capacity (Figure 2a, Appendix S1: Figure S1b) in the absence of trout. These galaxiid optima may be linked to the increasing proportion of unprotected consumers, such as mayflies and stoneflies, as food sources before overall prey biomass decreases at higher disturbance regimes (Jellyman and McIntosh 2020). Notably, the random effect of month also showed stronger effects for galaxiids than trout across all disturbance regime levels (Appendix S1: Figure S1b). These findings suggest that galaxiids are more adapted to seasonal changes, which might help these species coexist in dynamic environments. However, this adaptation could also make them even more vulnerable if climate change disrupts those seasonal patterns. (Hernández-Carrasco *et al*. 2025).

Supporting our model results, the proportion of locations with only trout in the observed data decreased as disturbance increased, while sites with only galaxiids showed the opposite trend (Fig. 3a). However, these observed results including densities (Figure 3b), overlook the underlying mechanisms that underpin community assembly, particularly species interactions and demography, with many of these co-occurrences likely population sinks for galaxiids (Woodford and McIntosh 2010). This mismatch between observed data and the underlying mechanisms highlights the risk of relying solely on descriptive statistics to inform conservation actions, predict invasion risks, and understand long-term trends (Tonkin *et al*. 2019). Our model performed strongly (Appendix 1: Figure S2, R^2^ = 0.73) and elucidates this issue, but it has limitations. Of particular note, our approach did not account for the movement of trout and galaxiids, particularly source-sink dynamics where galaxiids migrate from trout-free source populations to the studied locations. Strong propagule pressure from trout—especially in low disturbance regimes—may even lead to underestimates of the already strong impact of trout on native fish persistence across the natural disturbance gradient.

**Figure 3:**
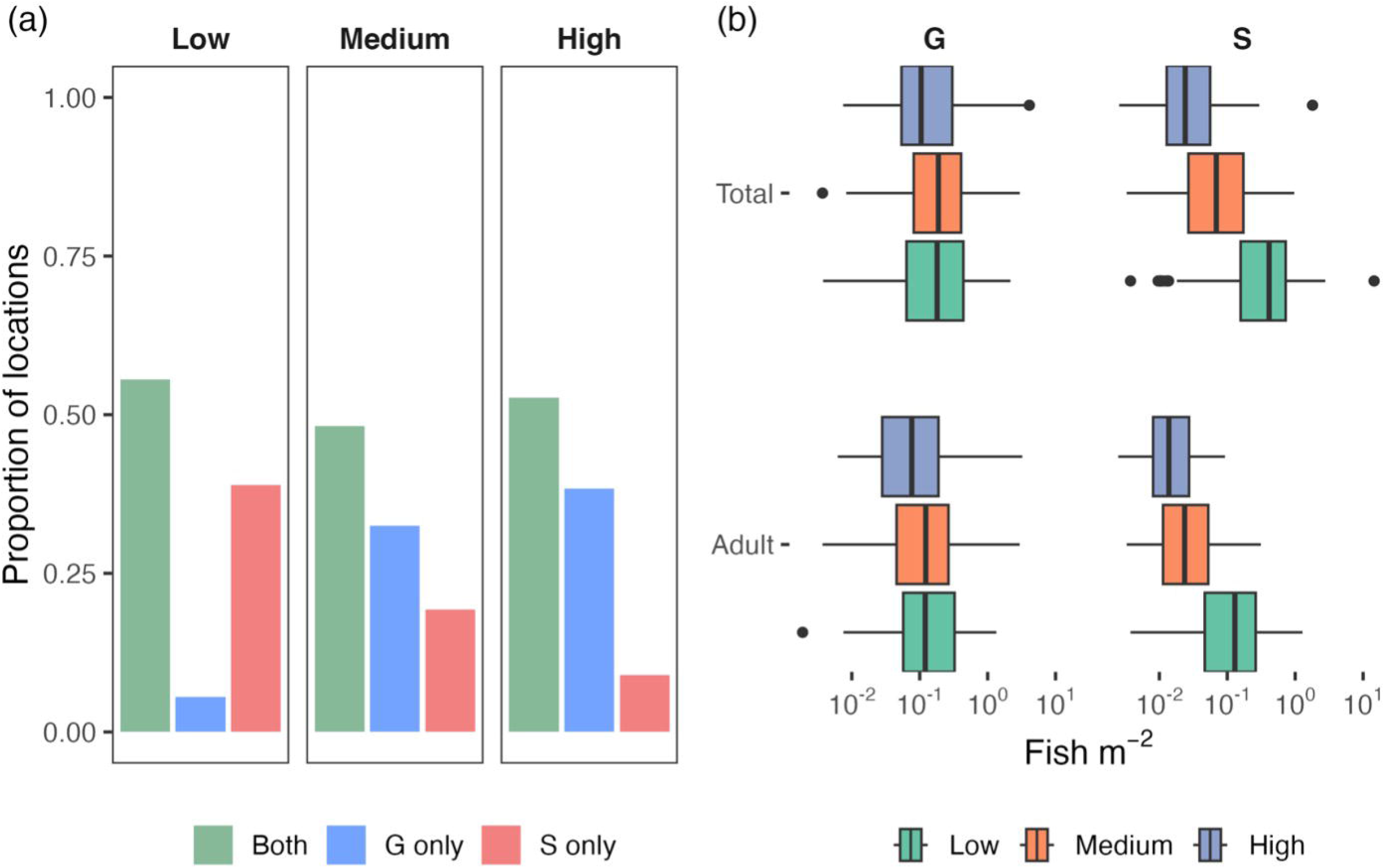
Observed densities of galaxiids (G) and trout (S) across the study area in the upper Waimakariri River catchment of New Zealand showing (a) how often fish are present at each level of disturbance, either exclusively or together with the competitor, and (b) the distribution of densities across locations, when present.

### Conclusion & management implications

Our results indicate that conservation strategies that take advantage of natural disturbance regimes to protect native species from invaders may not deliver the expected outcome. Here, native galaxiid fishes remained vulnerable to invasion by trout, even at high disturbance rates. Disturbance regimes structure competitive hierarchies and reveal signs of habitat partitioning (Stump and Vasseur 2023), but the clear imbalances favoring non-native trout dominance over native galaxiids across disturbance regimes found here make it clear that trout, like many other invaders, are an entrenched part of New Zealand’s freshwater ecosystems. As such, more targeted actions are needed to counter the extinction risk of native fish, such as explicit trout barriers and local removals (Jolly *et al*. 2024). Populations at sites with intermediate disturbance regimes, where our model predicts native fish to thrive best in the absence of trout, may be ideal focal points for conservation actions, including restoration actions targeting natural disturbance maintenance. But such actions require active biological interventions. Importantly, the high probability of priority effects at high disturbance regimes implies that harsh disturbance regimes are vulnerable to deliberate trout release or spillover from trout source populations, which could counteract conservation measures and tip those habitats towards a trout-dominated state. A misguided management with a sole focus on high disturbance regimes may turn out as an ecological trap (Battin 2004) when climate change and accompanying extreme events (Tonkin *et al*. 2019) increase harshness beyond what is tolerable.

