## Appendix S1 for "Disturbance provides limited respite for native fish against invaders"

### Supplement to:

#### Disturbance provides limited respite for native fish against invaders

Julian Merder[1], Angus R. McIntosh[1], Jan A. Freund[2], Rory S. Lennox[1], Naomi Heller[1], Jonathan D. Tonkin[1,3]

[1]School of Biological Sciences, University of Canterbury, Christchurch, New Zealand

[2]Institute for Chemistry and Biology of the Marine Environment, School of Mathematics and Science, Carl von Ossietzky Universität Oldenburg, Oldenburg, Germany

[3]Te Pūnaha Matatini Centre of Research Excellence, University of Canterbury, Christchurch, New Zealand

**Panel S1: Background on assumed population dynamics**

The two-species Lotka-Volterra competition model is given by (Chesson 2000):

$$\frac{1}{G_t}\frac{dG_t}{dt} = r_G(1 - a_{GG}G_t - a_{GS}S_t)$$
$$\frac{1}{S_t}\frac{dS_t}{dt} = r_S(1 - a_{SG}G_t - a_{SS}S_t)$$

where $G_t$ and $S_t$ represent the time dependent densities (fish/m$^2$) and $r_G$and $r_S$ maximal *per capita* growth rates of galaxiid (G) and trout (S) guilds, respectively. The competition coefficients $a_{GG}$ and $a_{SS}$ account for intraspecific competition, while $a_{GS}$ capturing interspecific competition effects. Integration over a time unit (one year), i.e., from $t$ to $t + 1$ , yields

$$ln\frac{G_{t+1}}{G_t} = r_G\left(1 - a_{GG}\int_t^{t+1} G_{t'}\,dt' - a_{GS}\int_t^{t+1} S_{t'}\,dt'\right) \approx r_G(1 - a_{GG}G_t - a_{GS}S_t)$$
$$ln\frac{S_{t+1}}{S_t} = r_S\left(1 - a_{SG}\int_t^{t+1} G_{t'}\,dt' - a_{SS}\int_t^{t+1} S_{t'}\,dt'\right) \approx r_S(1 - a_{SG}G_t - a_{SS}S_t)$$

where the approximation requires $G_t$and $S_t$ to match averages across seasons. Alternatively, the use of the rightmost term in the two equations above can be understood from the outset as a model formulation in discrete time. Since the extensive data set that we will use contains information about size-structured population strength for several sites and time points, but not necessarily for subsequent years $t$ and $t + 1$ we use the following surrogate method (exemplarily shown for G):
$G_t := G_{A,t} = G_{T,t} - G_{J,t}$ and $G_{t+1} := G_{A,t+1} = w_G G_{T,t} = w_G\left(G_{A,t} + G_{J,t}\right)$ where $G_{T,t}$ represents the total fish population in year $t$, split into adult $G_{A,t}$ and juvenile $G_{J,t}$ cohorts of the same year $t$ and $w_G$ is a survival factor (with value between 0 and 1) that carries adults and juveniles over to adults of the following year $t + 1$. In the following, we will suppress the temporal index $t$. Splitting the entire population into adults and juveniles is based on a threshold value, which is 65 mm for galaxiids and 100 mm for trout, guided by Boddy *et al.* 2020. Using this assumption, we have formulated a model as

$$ln(G_T) = -ln\, w_G + ln\, G_A + r_G(1 - a_{GG}G_A - a_{GS}S_A)$$

$$ln(S_T) = -ln\, w_S + ln\, S_A + r_S(1 - a_{SG}G_A - a_{SS}S_A)$$

whose parameters $(w_G, w_s, r_G, r_s, a_{GG}, a_{GS}, a_{SG}, a_{SS})$ can be fitted to age-structured abundance data $(G_A, G_J, S_A, S_J)$ from a single sampling time, with replicate samples taken at different locations.

For the growth rates, we used normal priors $\mathcal{N}(0.5, 0.5^2)$, truncated at 0 to ensure parameter estimates are positive. Similarly, the interspecific competition coefficients were assigned priors $\mathcal{N}(2, 2^2)$, also truncated at 0 to enforce non-negativity. The intraspecific competition coefficients were also assigned normal priors with the same mean and variance $\mathcal{N}(2, 2^2$, but with a stricter lower truncation at 0.5, based on the assumption that self-limitation must be sufficiently strong to prevent unsustainable densities (>2 fish/m²)—which are well above the expected carrying capacity, the inverse of this parameter. Although large trout can prey on galaxiids (Townsend & Crowl 1991), we assume that competition for habitat and resources is the dominant interaction, justifying a competition framework where all coefficients are strictly positive. For $w \in (0, 1)$, we assumed Beta priors, specifically Beta(5, 1), for S and G at each disturbance level, which concentrates most prior mass near one and decreases probability toward zero ($\sim 5^*w^4$). This reflects our belief that this term acts primarily as a correction, considering populations do not decline completely within a single year, and it avoids values near 0, making ln(w) approaching infinity unlikely. Seasonal differences in densities were modeled as a cyclic smooth function of month within the growth rate parameters to capture temporal heterogeneity, treated as a random effect in *brms* (Pedersen et al. 2019). For these smooth functions, we used horseshoe priors (df = 3), allowing seasonal adjustments to shrink toward zero if non-informative, thereby preventing overfitting (Piironen and Vehtari 2017). The residual standard deviations were modeled as $\mathcal{N}(0, 1^2)$ on the log link scale.

**Panel S3: Estimating competitive outcomes from modern coexistence theory**

Niche and fitness components are an essential part of modern coexistence theory. Coexistence between two species depends on their niche difference (ND) and fitness ratio (FR), following the framework of Chesson and Kuang 2008.
ND and FR are calculated as in Wan *et al*. 2024, with their values indicating the competitive outcome between G and S.

$$FR = \frac{F_S}{F_G} = \sqrt{\frac{a_{GG}a_{GS}}{a_{SS}a_{SG}}}$$

$$ND = 1 - \sqrt{\frac{a_{SG}a_{GS}}{a_{SS}a_{GG}}}$$

$$Coexistence: 1 - ND < FR < \frac{1}{1 - ND}$$

$$Priority\ Effect: 1 - ND > FR > \frac{1}{1 - ND}$$

If neither the coexistence nor the priority conditions are met, the FR decides about the competitive outcome, with FR > 1 indicating that S successfully invades and drives G to extinction.

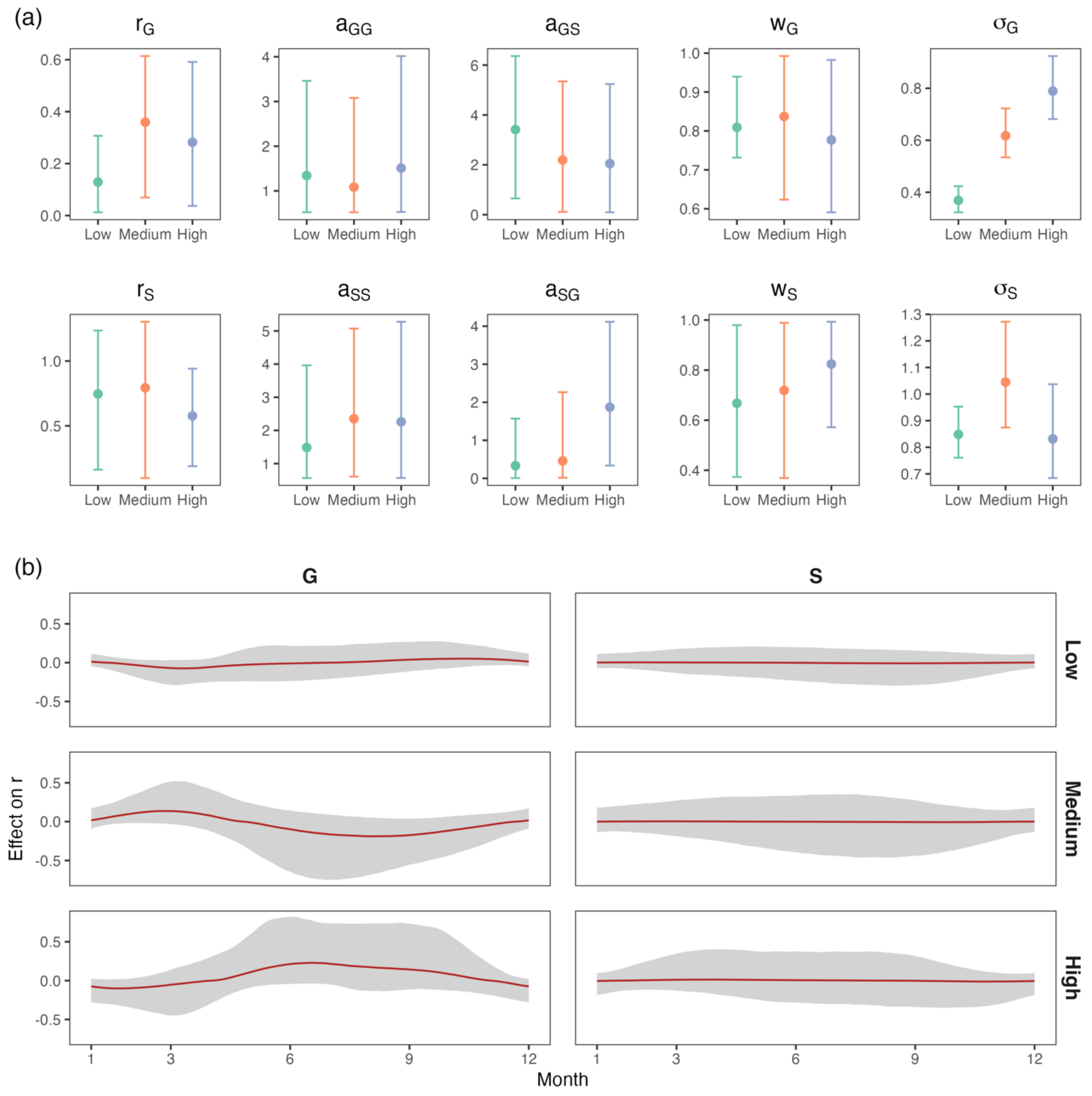


**Figure S1: Fitted model parameters for galaxiids (G) and trout (S).** Panel (a) indicates the fits of the fitted model parameters and standard deviations of log densities. Error bars indicate the 0.025-0.975 quantile range of parameters based on posterior draws from the model. Points represent the mean over posterior draws. Panel (b) shows the partial effect of month modelled as a random effect, thus seasonal deviation from the growth rate parameters for galaxiids and trout at each disturbance level shown in (a).

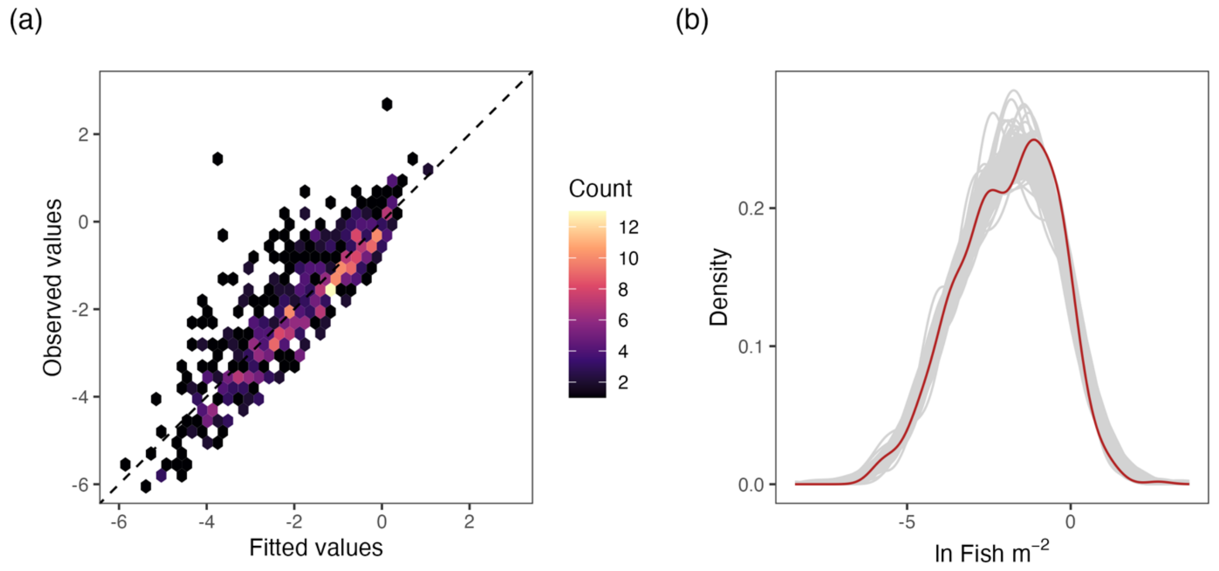


**Figure S2: Model validation plots.** Panel (a) shows fitted vs. observed values, where hexagonal bins are used to count points to avoid overplotting. Panel (b) shows the distribution of 100 posterior predictions from the model (grey), which aligns well with the observed distribution of fish densities (red).
